# Chronic adolescent stress increases exploratory behavior but does not change the acute stress response in adult male C57BL/6 mice

**DOI:** 10.1101/2021.06.25.449581

**Authors:** Oliver Sturman, Lukas von Ziegler, Mattia Privitera, Rebecca Waag, Sian Duss, Yannick Vermeiren, Peter de Deyn, Johannes Bohacek

## Abstract

Chronic stress exposure in adolescence can lead to a lasting change in stress responsiveness later in life and is associated with increased mental health issues in adulthood. Here we investigate whether the Chronic Social Instability (CSI) paradigm in mice influences the behavioural and molecular responses to novel acute stressors, and whether it alters physiological responses influenced by the noradrenergic system. Using large cohorts of mice, we show that CSI mice display a persistent increase in exploratory behaviors in the open field test alongside small but widespread transcriptional changes in the ventral hippocampus. However, both the transcriptomic and behavioural responses to novel acute stressors are indistinguishable between groups. In addition, the pupillometric response to a tail shock, known to be mediated by the noradrenergic system, remains unaltered in CSI mice. Ultra-high performance liquid chromatography analysis of monoaminergic neurotransmitter levels in the ventral hippocampus also shows no differences between control or CSI mice at baseline or in response to acute stress. We conclude that CSI exposure during adolescence leads to persistent changes in exploratory behavior and gene expression in the hippocampus, but it does not alter the response to acute stress challenges in adulthood and is unlikely to alter the function of the noradrenergic system.

## Introduction

Chronic stress is a major risk factor for neuropsychiatric disease, including anxiety and depression [1–4]. However, the majority of chronic stress research has focused on early life stress or adult stress exposure. Early life stress has been shown to lead to sustained hyperactivity and reactiveness of the hypothalamic-pituitary-adrenal (HPA) axis alongside hypertrophy of the adrenals, and dysregulation of glucocorticoid signalling in the hippocampus [5–10]. Chronic stress in adults generally leads to a similar dysregulation of the HPA axis/corticosterone levels and detrimental alterations to the hippocampus [8,11–13]. Meanwhile, adolescent/juvenile stress has received less attention. Still, it has been shown that chronic stress exposure during this sensitive developmental period can lead to alterations in the HPA axis [8,18,19] and the noradrenergic system [20–23]. Additionally, chronic stress exposure during adolescence can lead to enduring phenotypes that are different to those observed when stress exposure occurs during adulthood [14,15]. This is no surprise given the substantial reorganisation and the developmental processes occurring throughout the brain and HPA axis during this phase [16,17].

Adolescence is a period during which social environmental cues are particularly important [14–16]. The chronic social instability (CSI) paradigm [17] was created in order to investigate the effects of chronic stress exposure during adolescence. The paradigm involves randomly changing cagemates of mice twice a week for seven weeks in order to disrupt social hierarchies and introduce increased levels of social stress [18]. CSI leads to increased anxiety, increased corticosterone levels, hypertrophic and hyperresponsive adrenals, downregulation of both mineralocorticoid receptors and glucocorticoid receptors in the hippocampus and a flattening of the circadian rhythm [17,19–22]. Additionally it was shown that the anxiety behaviour may even persist transgenerationally, and that some of these effects can be reversed upon administration of selective serotonin reuptake inhibitors such as fluoxetine or paroxetine [17,22].

Ultimately, the goal of animal models in the study of chronic stress is to elucidate the neuroscientific mechanisms that underlie stress-related diseases. The stress response is complex, and goes beyond activation of the HPA axis [23,24]. Unbiased transcriptomic analyses have already highlighted several potential mechanisms responsible for the alterations in behaviour associated with chronic stress [25–31]. One key mechanism is a dysregulation of glutamatergic signalling pathways, which can lead to glucocorticoid-induced neurotoxicity and damage to the hippocampus [4,30,32]. Other studies have used transcriptomic screening to show that animals which are susceptible/resilient to stress show region-specific differences in gene expression, and have implicated brain derived neurotrophic factor (BDNF) and epigenetic changes as key mediators of the effects of chronic stress [28,33]. Transcriptomic profiling has also shown alterations in cortical plasticity following early life stress [34], and has pointed to altered mitochondrial gene expression profiles in the prefrontal cortex of susceptible mice [35]. Notably, metabolic alterations in high socially ranking mice are related to stress susceptibility [36,37]. These studies show that transcriptomic screens can provide valuable insights into the mechanisms at work behind chronic stress, and they suggest that social hierarchy is an important factor shaping stress resilience and susceptibility. However, to our knowledge, no studies have yet provided similar genome-wide datasets regarding chronic social stress exposure in adolescence.

Given previous studies link the noradrenergic system to the effects of chronic stress during adolescence [38–40], and, because noradrenaline plays a key role in the acute stress response and stress related disorders such as PTSD/depression/anxiety [41–43], we hypothesized that CSI would impact the noradrenergic system. To this end, we tested the noradrenaline dependent pupil response [44], and measured noradrenaline release in response to an acute stress challenge (swim) using ultra-high performance liquid chromatography (uHPLC). Since noradrenaline is a key regulator of the acute stress response, we also assessed behaviour following an acute stress cue (shock anxiety paradigm). Finally, we profiled the transcriptomic response to an acute restraint stress challenge after CSI treatment, and also performed a transcriptomic screen in the ventral hippocampus in a large cohort of CSI animals at baseline in adulthood.

## Methods

### Animals

C57BL/6J (C57BL/6JRj) mice (male, 2.5 months of age) were obtained from Janvier (France). Mice were maintained in a temperature- and humidity-controlled facility on a 12 hour reversed light–dark cycle (lights on at 08:15 am) with food and water ad libitum. Mice were housed in groups of 5 per-cage and used for experiments when 2.5-4 months old. For each experiment, mice of the same age were used in all experimental groups to rule out confounding effects of age. All tests were conducted during the animals’ active (dark) phase from 12-5 pm. Mice were single housed 24 hours before behavioral testing in order to standardize their environment and avoid disturbing cagemates during testing [45,46]. All procedures were carried out in accordance to Swiss cantonal regulations for animal experimentation and were approved under license 155/2015.

### Chronic Social Instability Paradigm (CSI)

We carried out the CSI procedure (based on the one developed by Haller and Schmidt [19,47] in two different cohorts of mice (n=60, n=59), each in a different building/lab. The mice arrived at the lab aged between postnatal day 21-23 housed in groups of 5. Upon arrival they were ear-tagged and split randomly (by cage) into either the CSI or control group. The CSI mice underwent the CSI paradigm, which consisted of briefly placing all CSI mice (n=30) into a larger cage, from which they were then randomly assigned to new cages. Mice in the control group were similarly handled, however, they entered the larger cage only with their cagemates before they were all returned to their original cage. The mice were subjected to these cage changes twice a week (Tuesday/Friday) for seven weeks, during the last cage change the mice were returned to their original cage and allowed to rest for 5 weeks prior to any further testing.

### Open Field Test (OFT)

Open-field testing took place inside sound insulated, ventilated multi-conditioning chambers (TSE Systems Ltd, Germany). The open field arena (45 cm x 45 cm x 40 cm [L x W x H]) consisted of four transparent Plexiglas walls and a light grey PVC floor. Animals were tested for 10 minutes under dim lighting (4 lux). Distance, time in center, supported rears and unsupported rears were recorded.

### Shock Anxiety

Shock Anxiety testing was performed as previously described [48] and took place inside the TSE multi conditioning systems’ black fear conditioning arena and the OFT arena described above. On day 1, all animals were placed into the fear conditioning arena for 3 minutes. After 2m 30s, animals in the shock group received a 3s 0.6mA shock. The following day, all animals were again placed into the fear conditioning chamber for 3 minutes before being directly placed into the OFT for 10 minutes.

### Pupillometry

Mice were anesthetized with 5% isoflurane before being transferred into a stereotactic frame and sustained with 2% isoflurane. Mice received two tail shocks (0.6mA for 3 seconds), the first, 2 minutes after being transferred to the frame and the second, 3:30mins after being placed in the frame. Recording was ceased after 5 minutes and the videos were then analysed using Deeplabcut as described previously [49]

### Forced Swim Test (FST)

Mice were forced to swim in a plastic beaker (20 cm diameter, 25 cm deep) filled to 17 cm with 17.9-18.1°C water for 6 minutes.

### Restraint Stress (RS)

Each mouse was restrained in a 50ml falcon tube, which had been modified so that the mouse could breathe through a small hole at the tip of the tube, whilst its tail could pass through a hole in the lid. The restraint tubes were placed back into the homecages, ensuring that they could not easily roll and that the breathing hole was not obstructed. Mice were removed from the restraint tubes after 30 minutes.

### Noldus EthoVision

EthoVision XT14 was used to acquire all forced swim and elevated plus maze videos and to analyse all of the open field videos. The automatic animal detection settings were used for all tests, slight tuning of these settings was performed using the fine-tuning slider in the automated animal detection settings to ensure the animals could be tracked throughout the entire arena. We ensured there was a smooth tracking curve and that the center point of the animal remained stable before analysis took place.

### DeepLabCut (DLC)

DeepLabCut 2.0.7 was used to track the performance of the animals in cohort 1 in the OFT. The data generated by DeepLabCut was processed using custom R Scripts that are available online (https://github.com/ETHZ-INS/DLCAnalyzer). For more information see [50].

### TSE Multi Conditioning System

Locomotion was tracked using an infrared beam grid; an additional beam grid was raised 6.5 cm above the locomotion grid to measure rearing. The central 50% (1012.5 cm^2^) was defined as the center of the arena. To automatically distinguish supported from unsupported rears, we empirically determined the area in which mice could not perform a supported rear [51]. Thus, all rears within 12.5 cm of the walls were considered supported rears, while rears in the rest of the field were considered unsupported rears. Rearing was defined as an interruption of a beam in the z-axis for a minimum of 150 ms. If another rear was reported within 150 ms of the initial rear, it was counted as part of the initial rear.

### Statistical Analysis of Behaviour

Data were tested for normality and all comparisons between normally distributed datasets containing two independent groups were performed using unpaired t-tests (2-tailed), data which was found to not follow a normal distribution was compared using non-parametric analyses. All comparisons between more than two groups were performed using two-way ANOVAs in order to identify group effects. Significant main effects were then followed up with post hoc tests (Tukey’s multiple comparison test).

### Tissue collection for uHPLC and immunohistochemistry

When brain tissue was collected for uHPLC, mice were rapidly euthanized by cervical dislocation. The hippocampus was immediately dissected on an ice-cold glass surface, snap-frozen in liquid nitrogen and stored at −80°C until further processing for uHPLC.

### Ultra-high performance liquid chromatography (uHPLC)

To quantify noradrenergic ((nor)adrenaline; 3-methoxy-4-hydroxyphenylglycol (MHPG)), dopaminergic (dopamine (DA); 3,4-dihydroxyphenylacetic acid (DOPAC); homovanillic acid (HVA)), and serotonergic (5-HT; 5-hydroxyindoleacetic acid (5-HIAA)) neurotransmitters and metabolites, a reversed-phase uHPLC system coupled with electrochemical detection (RP-uHPLC-ECD) was used (Alexys™ Neurotransmitter Analyzer, Antec Scientific, Zoeterwoude, Netherlands). In short, our previously validated RP-HPLC method with ion pairing chromatography was applied as described by Van Dam and colleagues [58; 59], albeit with minor modifications regarding the installed column (BEH C18 Waters column, 150 mm x 1mm, 1.7μm particle size) and pump preference (LC110S pump, 449 bar; isocratic flow rate of 58μL/min), achieving the most optimal separation conditions and a total runtime of under 15 minutes. Column temperature in the faraday oven was kept at 37°C. For electrochemical detection, a SenCell was used, with a glassy carbon working electrode (2mm) and an in-situ Ag/AgCl (ISAAC) reference electrode at a potential setting of 670mV and 5 nA output range. The mobile phase constituted a methanol-phosphate-citric acid buffer of 120mM citric acid, 120mM phosphoric acid, 600mg/L octane-1-sulfonic acid, 0.1mM EDTA, 8mM KCl, and, 11% methanol at pH 3.0. Levels of all monoamines and metabolites were calculated using Clarity software™ (DataApex Ltd., v8.1, 2018, Prague, Czech Republic).

Brain samples were defrosted to 4°C and subsequently homogenized in 800 μL ice-cold sample buffer, using a Minilys Personal homogenizer device and 1.4 mm ceramic zirconium oxide CK14 Precellys beads (Bertin Technologies SAS, France; 60s, speed rate 3)). Subsequently, to remove excess proteins, 450 μL homogenate was transferred onto a 10,000 Da Amicon® Ultra 0.5 Centrifugal Filter (Millipore, Ireland) that had been pre-washed once using 450 μL sample buffer (centrifugation: 14,000 × g, 20 min, 4°C). The Amicon® filter loaded with the homogenate was then centrifuged (14,000 × g, 20 min, 4°C). Finally, the filtrate was transferred into a polypropyelne vial (0.3mL, Machery-Nagel GmbH & Co. KG, Germany) and automatically injected (undiluted; 4x diluted with sample buffer) onto the previously-mentioned uHPLC column by the Alexys AS110 sample injector (4°C).

### Whole tissue RNA extraction

Samples were homogenized in 500 μL Trizol (Invitrogen 15596026) in a tissue lyser bead mill (Qiagen, Germany) at 4°C for 2 min at 20Hz, and RNA was extracted according to manufacturer’s recommendations. RNA purity and quantity were determined with a UV/V spectrophotometer (Nanodrop 1000), while RNA integrity was assessed with high sensitivity RNA screen tape on an Agilent Tape Station/Bioanalyzer, according to the manufacturer’s protocol.

### Sequencing

Library preparation and sequencing was performed at the Functional Genomics Center Zurich (FGCZ) core facility. For library preparation, the TruSeq stranded RNA kit (Illumina Inc.) was used according to the manufacturer’s protocol. The mRNA was purified by polyA selection, chemically fragmented and transcribed into cDNA before adapter ligation. Single-end sequencing (100nt) was performed with Illumina HiSeq 4000. Samples within experiments were each run on one or multiple lanes and demultiplexed. A sequencing depth of ∼20M reads per sample was used. Adapters were trimmed using cutadapt [52] with a maximum error rate of 0.05 and a minimum length of 15. Kallisto [53] was used for pseudoalignment of reads on the transcriptome level using the genecode.vM17 assembly with 30 bootstrap samples and an estimated fragment length of 200 ± 20. For differential gene expression (DGE) analysis we aggregated reads of protein coding transcripts and used R (v. 3.6.2) with the package “edgeR” (v 3.26.8) for analysis. A filter was used to remove genes with low expression prior to DGE analysis. EdgeR was then used to calculate the normalization factors (TMM method) and estimate the dispersion (by weighted likelihood empirical Bayes). We used a generalized linear model (GLM) with empirical Bayes quasi-likelihood F-tests. For multiple testing correction the Benjamini-Hochberg false discovery rate (FDR) method was used. For removal of artefacts and restraint effects from the CSI vs. Control comparison we further employed SVA correction to correct for processing specific effects [54]. Surrogate variables independent of experimental groups were identified using the sva package 3.34.0 on data after DESeq2 variance-stabilization [54,55], and were then included as additive terms in the GLMs.

## Results

### Cohort 1

We first carried out the CSI paradigm (n=59, cohort 1) to investigate the behavioural effects of adolescent stress exposure and to look for alterations in the noradrenergic system (Figure 1a). To ensure that any behavioral effects observed were not due to the random selection of mice, we performed an open field test (OFT) test prior to CSI exposure. As expected, we observed no significant differences between the behaviour of the mice in both groups (Supplementary Figure 1). Two weeks after completion of the CSI paradigm, we again tested mice in the OFT. We saw an anxiolytic effect of CSI, as CSI mice spent significantly more time in the center of the arena (unpaired t-test; t=3.033, df=57, p=0.0036) (Figure 1c). The CSI mice in this cohort also covered significantly more total distance (unpaired t-test; t=4.395, df=57, p<0.0001) (Figure 1b). The observation that CSI treatment leads to increased exploration was unexpected, as previous reports in adolescent CD1, and adult C57BL6/J mice had suggested an increase in anxiety related measures after CSI exposure [19,20,22]. In the light-dark-box test one week after the OFT we observed no differences between groups on any of the measures (See Figure 3f-i). This suggests that CSI increases exploratory behavior in adolescent male C57Bl/6 mice in a context-dependent manner.

**Figure 1.**
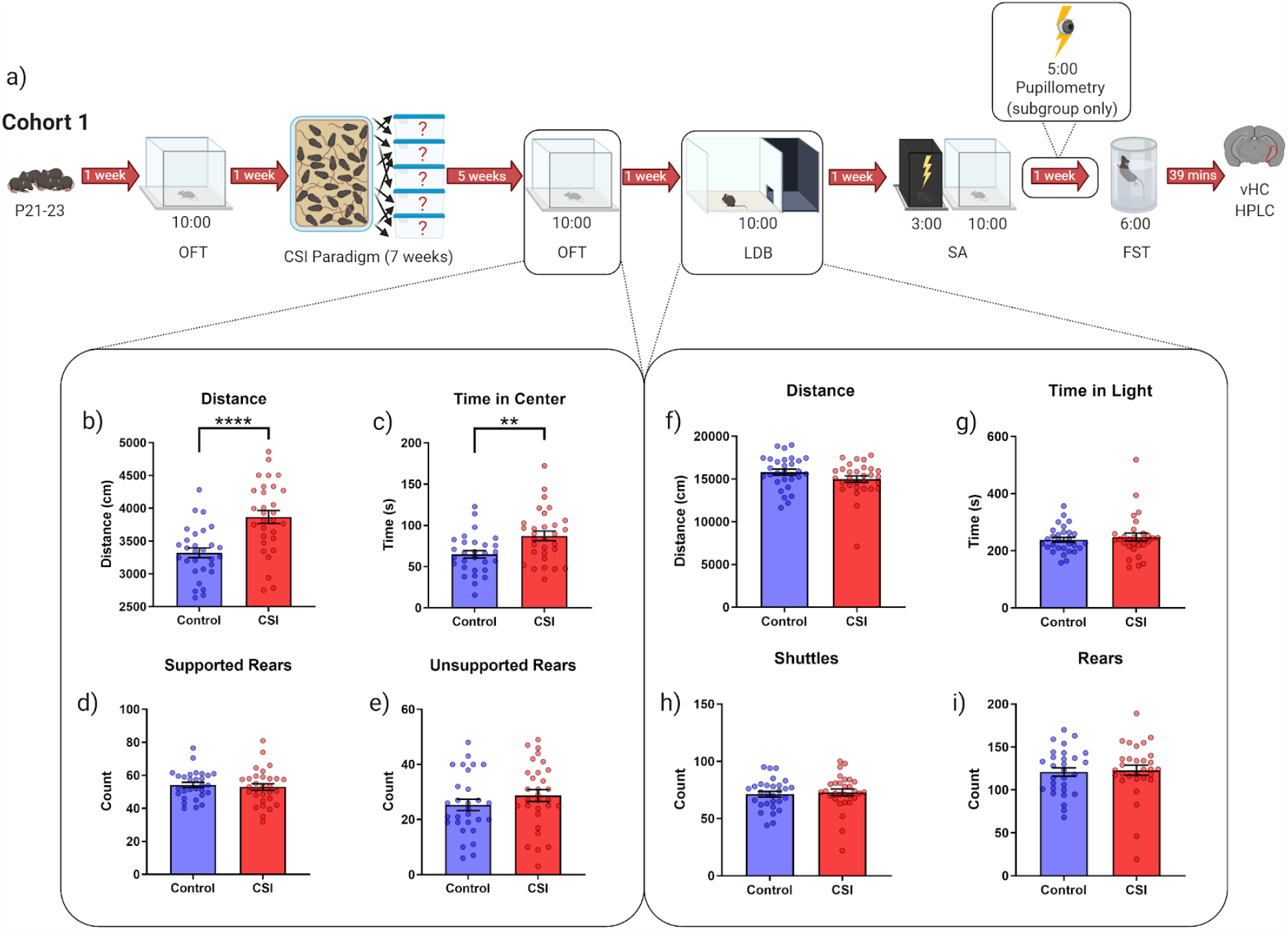
Persistent behavioral changes induced by CSI stress. (a) schematic outline of the experimental procedures performed on cohort 1. (b-e) Distance, time in center, supported and unsupported rears in the open field test as reported by our custom DeepLabCut analysis suite (see methods).(f-i) Distance, time in light, shuttles and rears in the light dark box as reported by the TSE Multi Conditioning System. Data expressed as mean ± SEM. **=p<0.01,****=p<0.0001.

### Shock Anxiety/Acute Stress following CSI

To further investigate the behavioural response of the CSI animals to a novel acute stressor, mice from cohort 1 were subjected to a shock anxiety paradigm in which animals were placed into a fear conditioning chamber, where half of the animals received a shock (shock group), the other half did not (control group). The following day, all animals were briefly placed back into the shock box (but this time none of the mice received a footshock). This served as a brief anxiety cue for the shock group [48]. Immediately afterwards, mice were placed into an open field (see Figure 2b/methods). It appears that the memory of the animals was intact given the significant main effect of shock on distance (Two-Way ANOVA, F(1,55)=22.48, p<0.0001) and total number of rears (Two-Way ANOVA, F(1,55)=29.3, p<0.0001) when placed into the shock context prior to the OFT on day 2 of the paradigm (Supplementary Figure 2). The effects of this anxiety cue persisted in the following open field test, as we observed a significant main effect of shock on distance (Two-Way ANOVA, F(1,55)=13, p=0.0007) and supported rears (Two-Way ANOVA, F(1,55)=5.754, p=0.0199) (Figure 2c,e). However, no significant interactions were observed, implying that the control and CSI animals responded similarly to the novel stressor in this paradigm.

**Figure 2.**
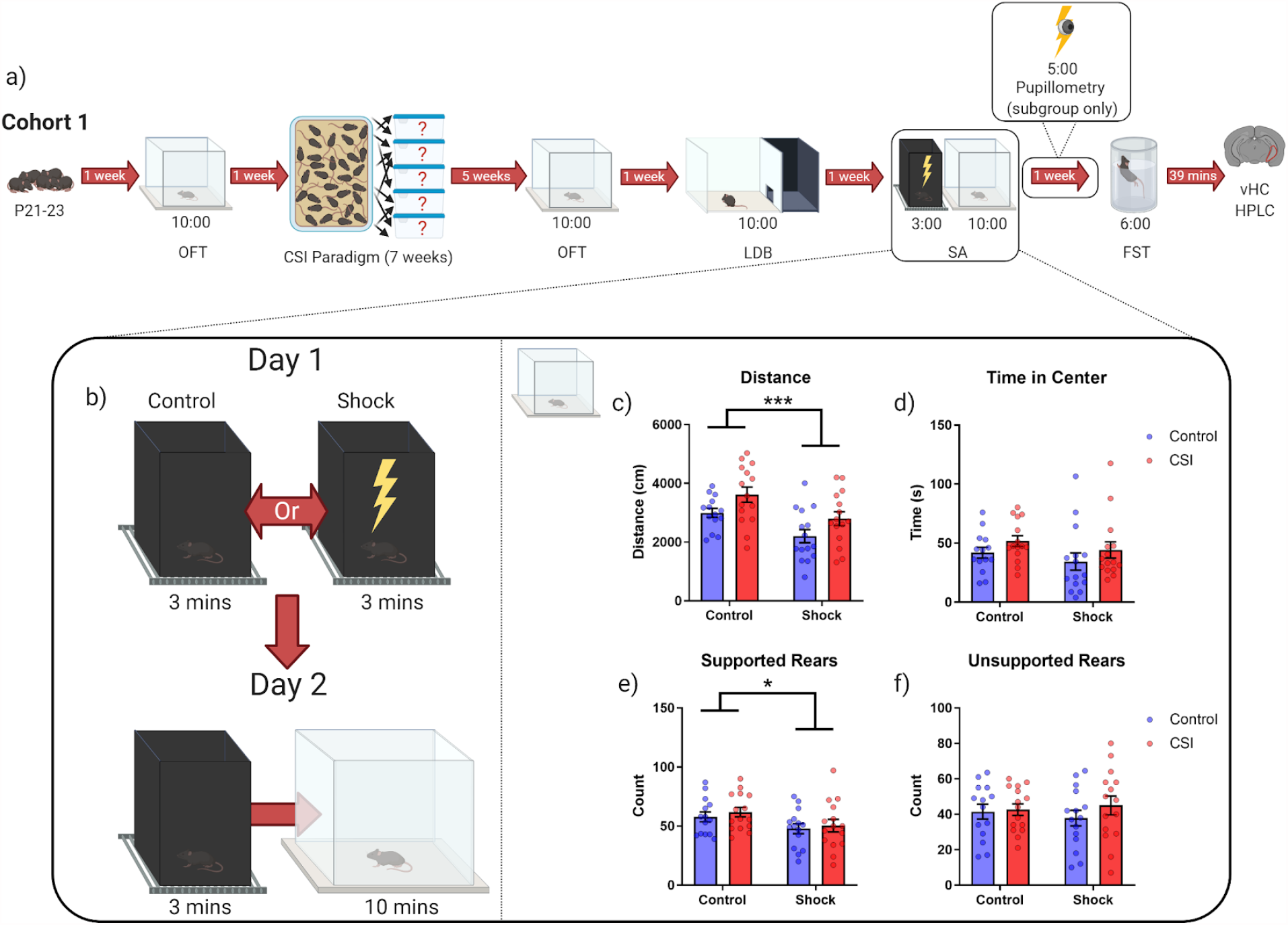
CSI and control mice in a shock-anxiety paradigm. (a) schematic outline of the experimental procedures performed on cohort 1. (b) detailed schematic shock anxiety protocol. (c-f) Distance, time in center, supported rears and unsupported rears as reported by our custom DeepLabCut analysis suite (see methods) in the OFT immediately following brief re-exposure to the shock context. Data expressed as mean ± SEM. *=p<0.05,***=p<0.001.

**Figure 3.**
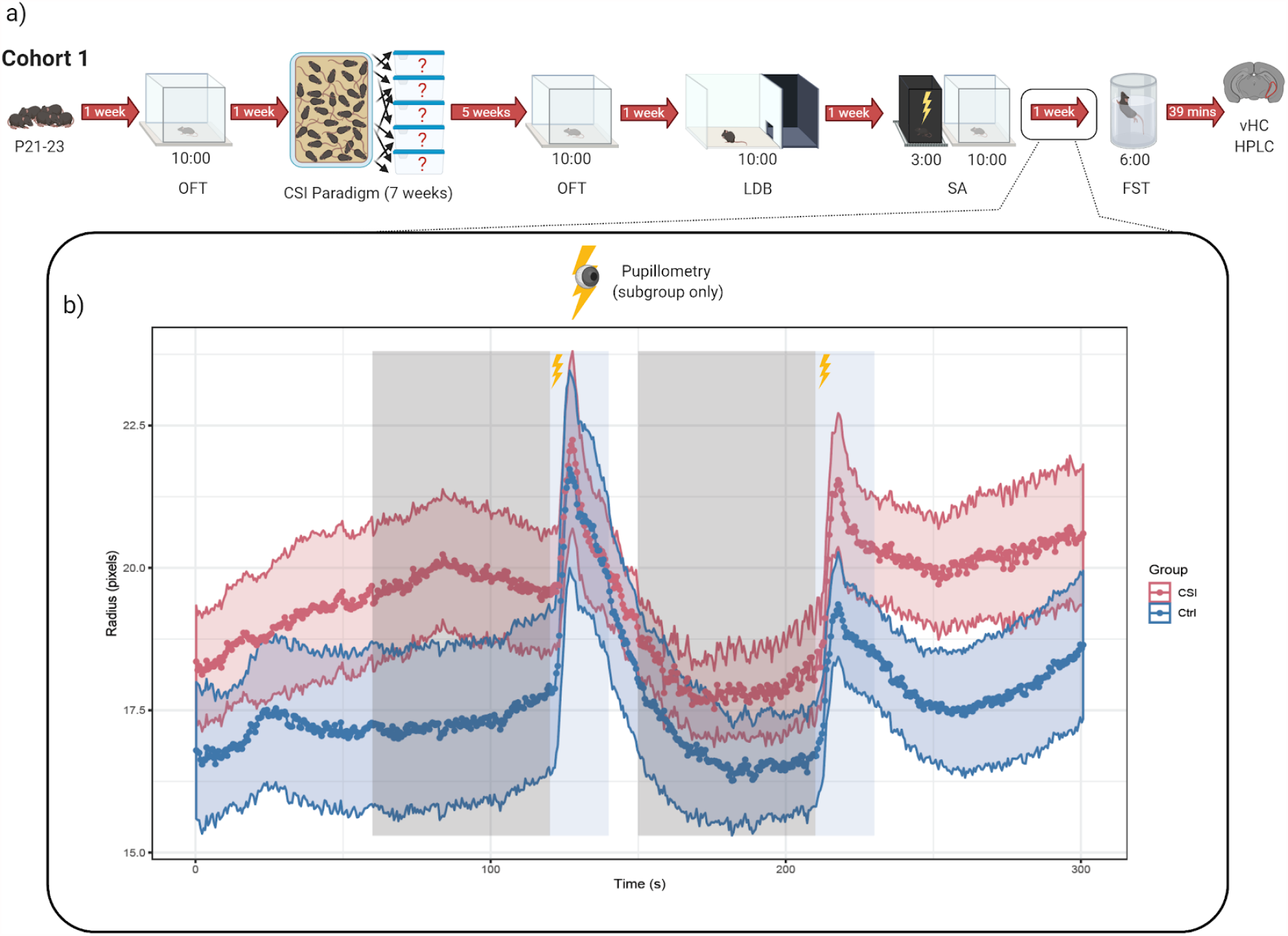
Shock-induced pupillometry in CSI and control mice. (a) schematic outline of the experimental procedures performed on cohort 1. (b) pupil radius of CSI (n=14) and control mice (n=15). Shock responses are indicated in blue, with the corresponding baseline pupil radii indicated in the grey, data expressed as mean ± SEM.

### Pupillometry

Chronic stress can also induce persistent neuroplastic alterations to the locus coeruleus - noradrenergic system [56,57]. Therefore, we wanted to see if the CSI paradigm had altered the pupil response of the animals, which is known to be regulated by the locus coeruleus-noradrenergic system and can be measured in response to a tail shock under light isofluorane anesthesia [49]. We therefore examined the pupillometric response of a subset of the mice to a tail shock. We selected those mice which had received a shock in the shock anxiety test, (n=14 CSI, 1 mouse excluded due to equipment malfunction), n=15 control). Although the electrical stimulus elicited a measurable dilation of the pupil (linear mixed effects model (’Radius∼Group*Response+(1|Animal)+Time’ against ’Radius∼(1|Animal)+Time)β =1.5933, SE=0.7899, t(84)=2.017, p=0.0469), we observed no significant differences between control and CSI animals at baseline (β =-1.8315, SE=1.3808, t(37.6730)=-1.326, p =0.1927)) and no significant group/response interaction (β =0.7140, SE=1.0984, t(84) =0.650, p=0.5174)(Figure 3b). These results suggest that the pupil response to a novel stressor is not altered by a history of CSI exposure.

### Noradrenaline turnover

Since pupil dilation is an indirect measure of locus coeruleus activation, we decided to directly measure noradrenaline release and turnover in the vHC (a direct projection region of the locus coeruleus) after an acute forced swim stress exposure. To this end, we used uHPLC, which can measure several monoamines and their metabolites simultaneously with high precision [58,59]. We had shown previously that noradrenaline turnover is a sensitive readout of locus coeruleus activation [49,60]. In both CSI and control mice, we detect strong main effects of acute stress, with significant increases in dopamine (DA, Two-Way ANOVA, F(1,26)=51.45, p<0.0001), the noradrenaline metabolite 3-methoxy-4-hydroxyphenylglycol (MHPG, Two-Way ANOVA, F(1,26)=56.83, p<0.0001), the dopamine metabolite homovanillic acid (HVA, Two-Way ANOVA, F(1,26)=86.03,p<0.0001), and the serotonin metabolite 5-hydroxyindoleacetic Acid (5HIAA, Two-Way ANOVA, F(1,26)=8.594, p=0069). Additionally, noradrenaline turnover was strongly increased (MHPG/NA ratio;Two-Way ANOVA, F(1,26)=36.18, p<0.0001), dopamine turnover was suppressed (HVA/DA ratio; Two-Way ANOVA, F(1,26)=5.215, p=0.0308) and serotonin turnover was increased (5-HIAA/5-HT ratio; Two-Way ANOVA, F(1,26)=6.914, p=0.0142) (Figure 4d-l). However, we detected no interaction between treatment (acute stress) and group (control/CSI) for any of the measurements. In addition, we also analyzed the total distance travelled in the forced swim test and the time spent floating, as this has been used as a measure of monoaminergic activity and stress-responsiveness in the past [61–63]). We observed no differences between CSI and control mice in distance (unpaired t-test; t=0.7183, df=27, p=0.4787) or time floating (unpaired t-test; t=0.5579, df=27, p=0.5815). Again, these findings support the overall conclusion that the effects of acute stress are not altered in animals with a history of CSI exposure (Figure 4b,c).

**Figure 4.**
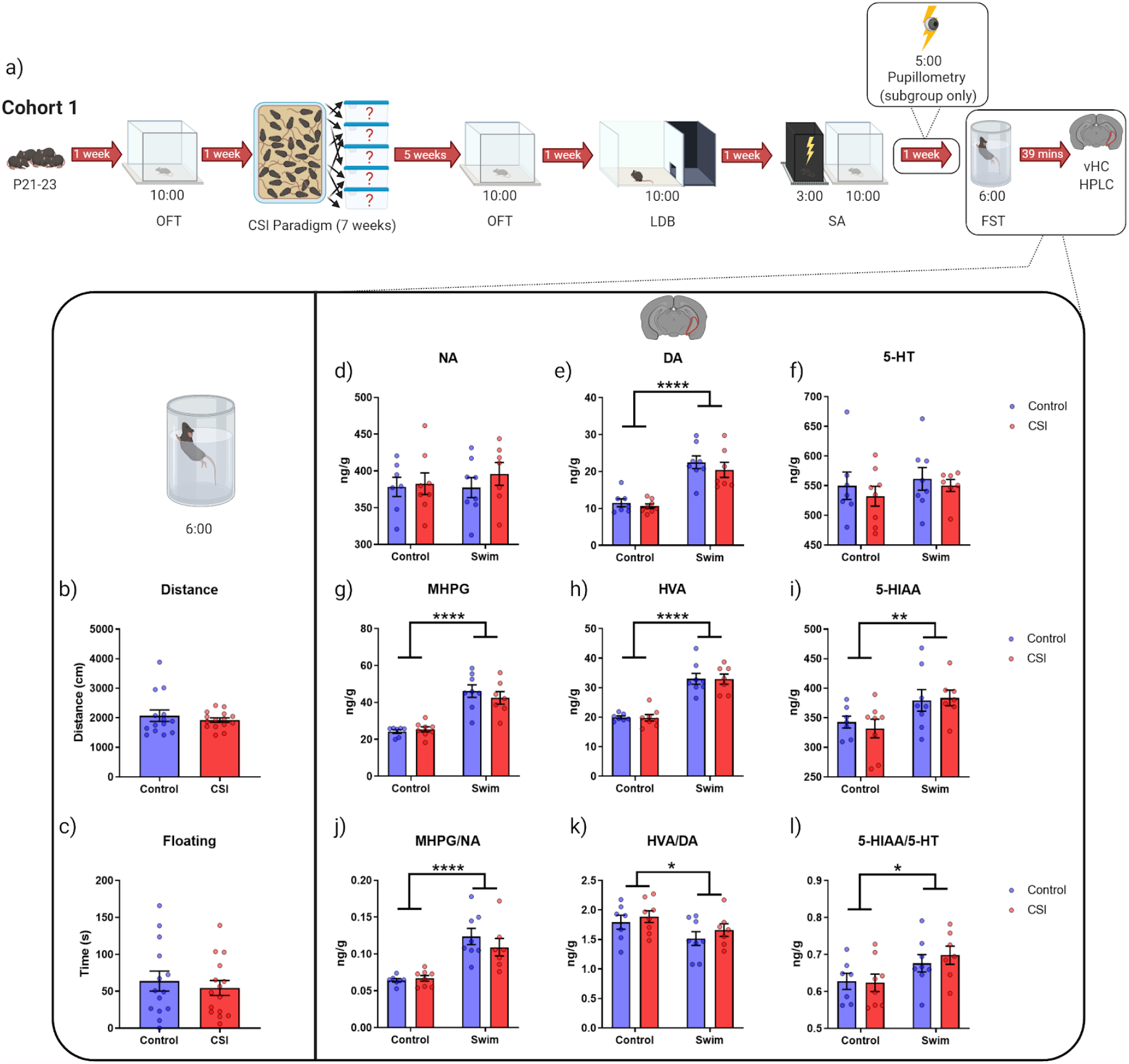
Stress-induced changes in monoaminergic neurotransmitter systems in CSI and control mice. (a) schematic outline of the experimental procedures performed on cohort 1. (b-c) Distance/time floating in the FST as reported by Noldus Ethovision XT13.(d-l) uHPLC analysis of: noradrenaline (NA), dopamine (DA), serotonin (5-HT), 3-methoxy-4-hydroxyphenylglycol (MHPG), homovanillic acid (HVA), 5-hydroxyindoleacetic acid (5HIAA), MHPG/NA, HVA/DA and 5-HIAA/5-HT in the ventral hippocampus post swim stress. Data expressed as mean ± SEM. *=p<0.05,***=p<0.001.

### Cohort 2 - transcriptomic changes induced by chronic social instability stress

#### Behavioural Differences

Given that we did not find support for our hypothesis that the acute stress response was altered after a history of CSI exposure, we decided to perform a transcriptomic screen to assess whether on a transcriptome-wide level we could detect differences in the acute stress response between CSI and control mice. Because transcriptomic analyses are extremely sensitive to the impact of acute or chronic stressors[25,34,64], we hypothesized that prior exposure to chronic stress would alter the subsequent transcriptomic response to acute stress. We generated a new cohort of CSI mice (n=30 controls, n=30 CSI, cohort 2). To show that CSI was effective, we again performed an OFT 5 weeks after the end of the CSI paradigm. Similar to cohort 1, we again saw an increase in exploratory behaviors after CSI treatment. CSI mice spent significantly more time in the center of the arena (Mann Whitney test; U=289, Sum of ranks=754(a) 1076(b), p=0.0167) (**Figure 5c**) and performed significantly more supported (unpaired t-test; t=2.215, df=58, p=0.0307) and unsupported rears (unpaired t-test; t=3.636, df=58, p=0.0006) (**Figure 5d-e**). Due to a physical relocation of the lab, all experiments and procedures carried out in this cohort took place in different animal housing and behavior testing facilities. Differences in behavioural recording apparatus (camera angle) meant that we were unable to use our deep-learning based analysis on this cohort [50]. However, experiments were carried out under comparable testing conditions (same mouse strain and vendor, same experimenter, same lighting conditions, same testing apparatus, same experimental procedures). We were able to analyze cohort 1 using the TSE system, thus we were able to pool data from cohort 1 and cohort 2. This resulted in a very large group size (n=60 CSI, n=59 control) and the analyses confirmed that overall CSI mice spent more time in the center of the open field (Mann Whitney test; U=1096, Sum of ranks=2866(a) 4275(b), p=0.0003) and performed significantly more unsupported rears (Mann Whitney test; U=1179, Sum of ranks=2949(a) 4191(b), p=0.0015). Combined, this suggests an increase in exploratory behaviors and a reduction in anxiety following CSI treatment (Supplementary Figure 3). Since others using the CSI paradigm have reported differences in locomotor adaptation [20], we also investigated this parameter in both cohorts but found no significant differences between control or CSI mice (data not shown).

**Figure 5.**
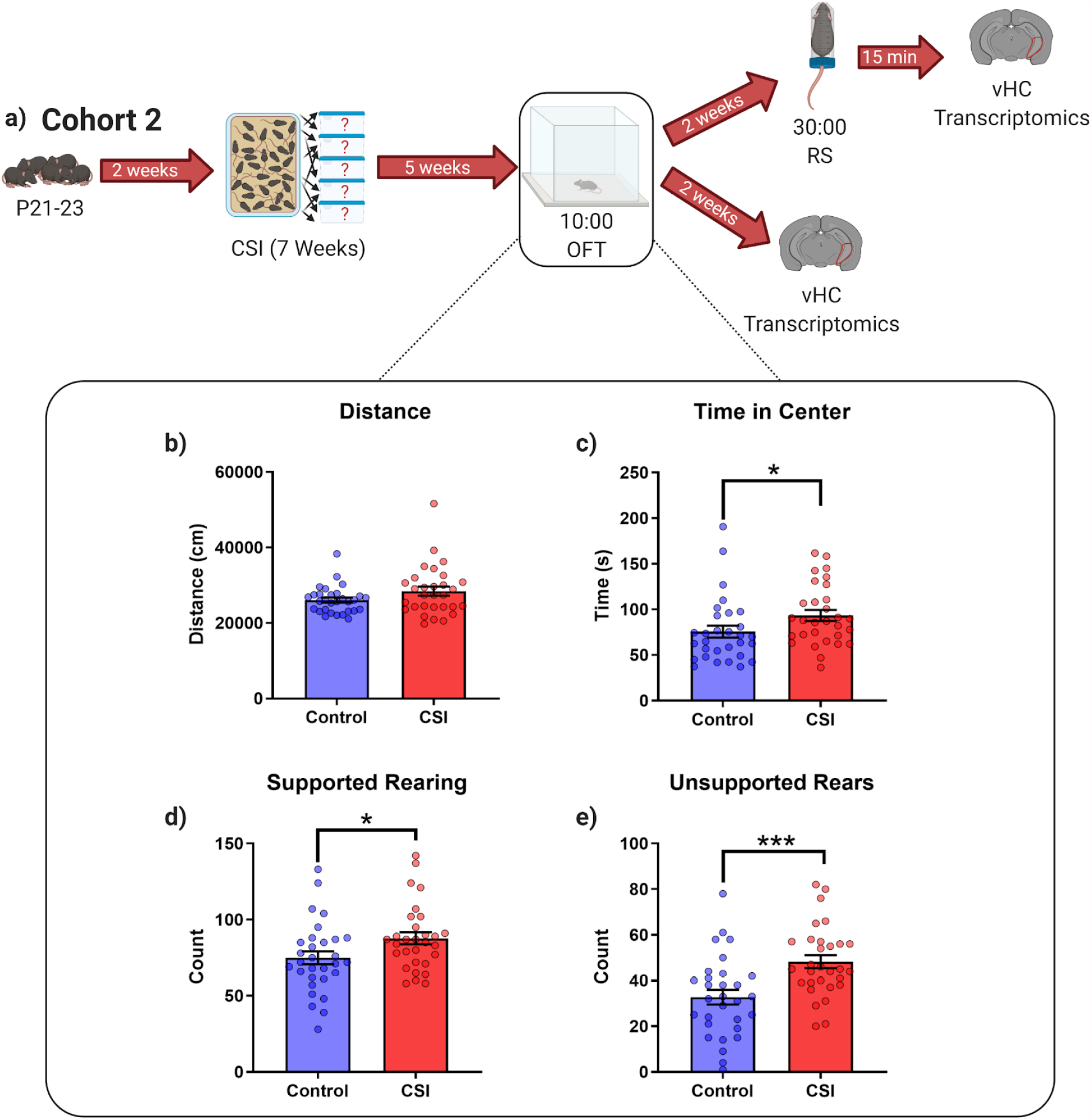
Long-lasting behavioral effects of CSI stress in a second cohort of mice. (a) schematic outline of the experimental procedures performed on cohort 2 (b-e) Distance, time in center, supported and unsupported rears in the open field test as reported by the TSE Multi Conditioning system. Data expressed as mean ± SEM. *=p<0.05, **=p<0.01,***=p<0.001.

#### Transcriptomic Differences

Two weeks after behavior testing, we performed transcriptomic profiling of the ventral hippocampus (vHC), a brain area known to be highly sensitive to the effects of stress [28,64,65]. To avoid selection bias, we decided to sequence the vHC of each mouse. The majority of mice were euthanised for tissue collection directly from their homecage (n=24 CSI; n=24 control), while a small cohort was exposed to 30min acute restraint stress, starting 45min before tissue collection (n=6 CSI, n=6 control). Without acute stress exposure, we detected many transcripts that were significantly up- (1748 genes) or down-regulated (1997 genes) between the control and CSI groups (**Figure 6a, Supplementary table 1**). We observed considerable co-expression and co-variability of many of the significant genes. To ensure that these observations were not artefacts we used surrogate variable analysis (SVA) to remove technical variability across batches of samples before re-analysing for CSI effects [54]. We were able to detect 377 robustly altered transcripts (73 up-regulated and 304 down-regulated) (**Figure 6d, Supplementary table 2**), confirming that CSI induces widespread changes of modest effect size. In response to acute restraint stress, we detect the expected strong transcriptomic differences we had already described previously [64,66] (**Figure 6a, b, Supplementary table 3**). These include predominantly well-described transcription factors known to be strongly activated by various stressors (**Figure 6b**). Contrary to our hypothesis, we detected no significant interaction between acute stress (restraint) and chronic stress (CSI) (**Figure 6a, Supplementary table 4**). Thus, prior exposure to CSI stress does not alter the strong, stereotypical transcriptional response to acute stress in the vHC. This observation is in line with the behavioural data showing no significant interaction between control and CSI animals following acute stress. Gene ontology enrichment analysis on the genes significantly regulated by acute restraint stress highlights the impact of the restraint stress on pathways related to memory formation and energy metabolism (**Figure 6c**), which is in line with our current understanding that the acute stress response mobilizes energy substrates and regulates energy homeostasis [24]. Gene ontology enrichment analysis of the large number of significant genes altered by CSI treatment reveals an interesting suppression of pathways related to synaptic function and glutamatergic signaling (Figure 6e). Although this reduction is rather subtle for each individual gene (5-10%), the convergence of these genes on key players of synaptic function (e.g. Shank1, Shank3, Grin1, Grin2d, Dlgap3, Dlgap4) is striking (**Figure 6e**).

**Figure 6.**
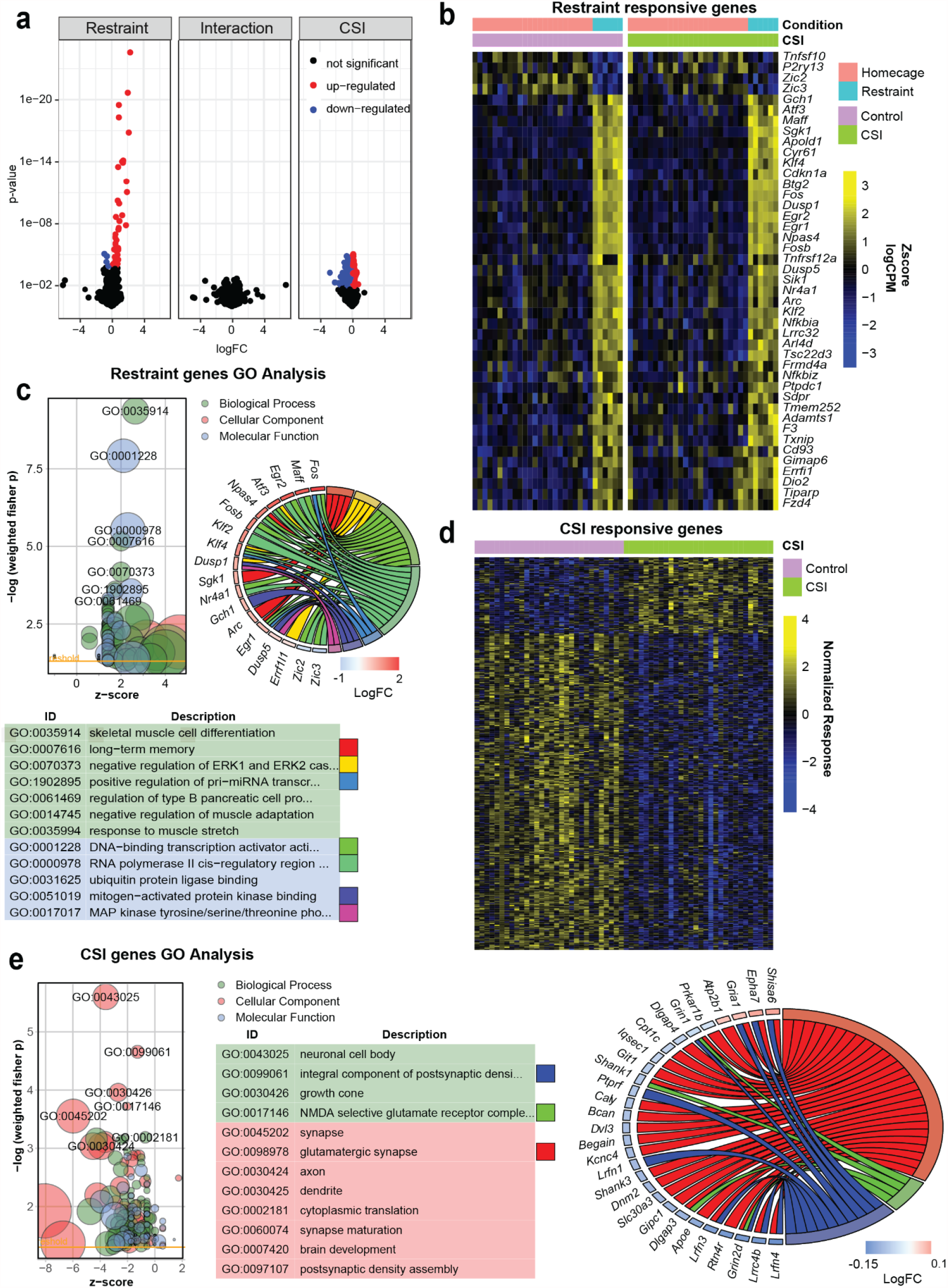
Transcriptomic changes in CSI and control mice at baseline and after acute restraint stress. **a)** Volcano plots visualizing log fold changes vs p-values for coefficients of an interactive model (CSI * restraint). Colors indicate significantly (adj.p < 0.05) up- (red) or down-regulated (blue) genes. **b)** Heatmap of genes altered by restraint stress in CSI and control animals across all samples **c**,**e)** Functional annotation results for significant restraint genes **(c)** or CSI genes (**d)** Heatmap of genes altered by CSI across all samples after removal of variability. **(e)**. *Box:* z-score vs significance of GO terms. Bubble size indicates number of assigned genes. Negative z-scores indicate that these genes are down-regulated, positive that they are up-regulated. Colors indicate the GO category. *Table:* top enriched (-log10(weighted fisher p) > 2.5 for restraint, > 3 for CSI) GO terms and descriptions. Boxes on the right indicate colors of these terms in the circular plot. *Circle:* GO chord plot illustrating gene to GO term associations for selected terms. Differentially regulated genes are indicated on the left side, GO terms on the right side. GO terms colors correspond to the legend in the GO table. A connection between GO term and gene indicates that the gene is part of the GO term. Color of the small box next to the genes indicates their logFC after restraint **(c)** or CSI **(e)**.

## Discussion

Here we show that chronic social instability (CSI) stress leads to an increase in the time spent in the center of the OFT alongside an increase in unsupported rearing behaviour 5 weeks after the end of the CSI paradigm. This anxiolytic, pro-exploratory phenotype is accompanied by widespread yet subtle transcriptomic alterations in the ventral hippocampus between control and CSI mice. However, we did not see any further behavioural differences between CSI/control mice in the LDB or following exposure to the shock anxiety paradigm or in the forced swim test. In addition, we saw no physiological differences in the pupil response following tailshock or in the neurotransmitter levels during uHPLC analysis of the ventral hippocampus at baseline or post swim stress following CSI. On the whole, our data do not support the hypothesis that a history of CSI alters the subsequent response to an acute stressor, nor alters the functional release of NA/MHPG or related monoaminergic neurotransmitters in the vHC, which is directly linked to the locus coeruleus.

The literature on the behavioral effects induced by adolescent CSI exposure is conflicting. Some studies have reported increased anxiety levels as indicated by decreased locomotor adaptation in the OFT, increased latency to eat in the novelty suppressed feeding test, and fewer open arm entries in the elevated plus maze [17,19–21]. However, these experiments were conducted in outbred CD1 mice, during the light phase of the light-dark cycle, and varied between males and females. Another group exposed adult CD1 mice to the CSI paradigm, tested them during the dark cycle and found no differences in anxiety-related behaviors, but instead an increase in rearing in response to CSI exposure [67,68]. Since we repeated the paradigm in two separate cohorts of animals in two entirely different facilities, we are confident that the behavioural phenotype we observed is reproducible under the testing conditions used in our lab/labs. Despite the reproducible effect of CSI on OFT behavior, it was surprising that we saw no differences in the LDB, where one would expect a decrease in anxiety would lead to an increase in time spent in the light compartment, however there is often poor agreement between different anxiety tests [69,70]. Given the lack of a detectable difference between control and CSI animals following novel acute stressors, it could be that the increased exploratory drive observed in the OFT indicates that CSI mice enter minimally stressful novel environments with higher levels of curiosity, but appear to be just as susceptible to the effects of more severe stressors (or more anxiogenic environments) as control mice. Indeed, the stress response of CSI mice is indistinguishable from controls in terms of transcriptomics, shock anxiety response, pupillometry, and uHPLC analysis of monoaminergic neurotransmitters in the ventral hippocampus. It is possible that the acute stress response elicited by the novel stressors (shock, cold swim) is too ‘hard wired’ to be altered by a chronic stress paradigm of this variety and intensity, or that the length of time between the end of chronic social instability and some of the final behavioural tests (>7 weeks) was too long, giving the animals enough time to recover from the stressor.

Although the noradrenergic system has been implicated in behavioural changes after stress exposure [38–40,56], and, linked to depressive-like behaviours following chronic stress [71,72], we were unable to show alterations in monoaminergic content in mice which show altered behaviour following adolescent social stress. This could be due to our use of an indirect measure of global noradrenergic activity (pupillometry), and the fact that our molecular analyses (uHPLC/transcriptomics) were restricted to one brain region (ventral hippocampus). Directly assessing the activation of noradrenergic brain regions such as the locus coeruleus during behaviour with electrophysiology, calcium imaging or voltammetry may show more transient alterations in the monoaminergic system that are overlooked here. However, it appears more likely that social stress during adolescence does not elicit its persistent phenotype in mice via alterations to monoaminergic transmission, as has been shown by others in rats [73].

Our transcriptomic analysis of the ventral hippocampus of CSI animals at baseline implies that NMDA receptors and differences in synaptic maturation and development may be responsible for the observed baseline differences between CSI mice and controls that led to the OFT phenotype (Figure 2C). In addition, we also find several genes (EPHA7,GRIN1,GRIN2D,PRKAR1B and SLC30A3) under the GO term: ‘hippocampal mossy fiber to CA3 synapses’ to be downregulated in CSI mice. Interestingly, there is a link between the hippocampus, mossy fibers and rearing in the context of novelty. Genetic selection for increased novelty-induced rearing leads to larger intra/infra pyramidal-mossy fiber terminal fields and potentially more efficient spatial processing [74–76]. It would therefore be interesting to test whether CSI treatment affects mossy-fiber structure and function.

Following exposure to acute restraint stress, both CSI and control mice exhibit a similar transcriptional response with GO enrichment analysis highlighting responses to the metabolic cost of enduring a stressor. These results are in line with the highly metabolically demanding processes triggered by stress, such as increased synaptic activity and increased glutamate release throughout the brain, specifically in stress sensitive regions such as the hippocampus [32,77,78]. However, what we see in the CSI mice could be a subtle yet meaningful downregulation of excitatory glutamatergic pathways, possibly to counteract exocytoxicity from sustained glutamate levels. NMDA selective glutamate receptors are downregulated in CSI mice, and it has been shown that blocking NMDA receptors and lowering the excitatory activity of ion channels leads to reductions in stress-induced-dendritic-remodelling in the hippocampus, preventing the maladaptive effects of stress [79,80]. In line with our hypothesis, this should be investigated in more depth e.g. using electrophysiology, before drawing further conclusions.

## Acknowledgements

We would like to thank animal caretakers Jens Weissman, Roger Staub and Sanja Vasic for looking after our animals, the lab of Isabelle Mansuy for allowing us to use her lab space and animal facility during the first year of this project, Han-Yu Lin for sample preparation and Marcus Grüschow for stimulating discussions.

## Data availability

All sequencing data are available to readers via Gene Expression Omnibus with identifier GSE172451 https://www.ncbi.nlm.nih.gov/geo/query/acc.cgi?acc=GSE172451

## Author Contributions

Conceptualization, OS, LvZ, JB; Methodology, OS, LvZ, MP, YV, PdD ; Investigation, OS, RW, SD, MP, YV, PdD ; Writing—Original Draft, OS, LvZ, JB; Writing—Review & Editing, OS, LvZ, YV, PdD, JB; Funding Acquisition, JB; Resources, JB.

## SUPPLEMENTARY FIGURES

**Supplementary Figure 1.**
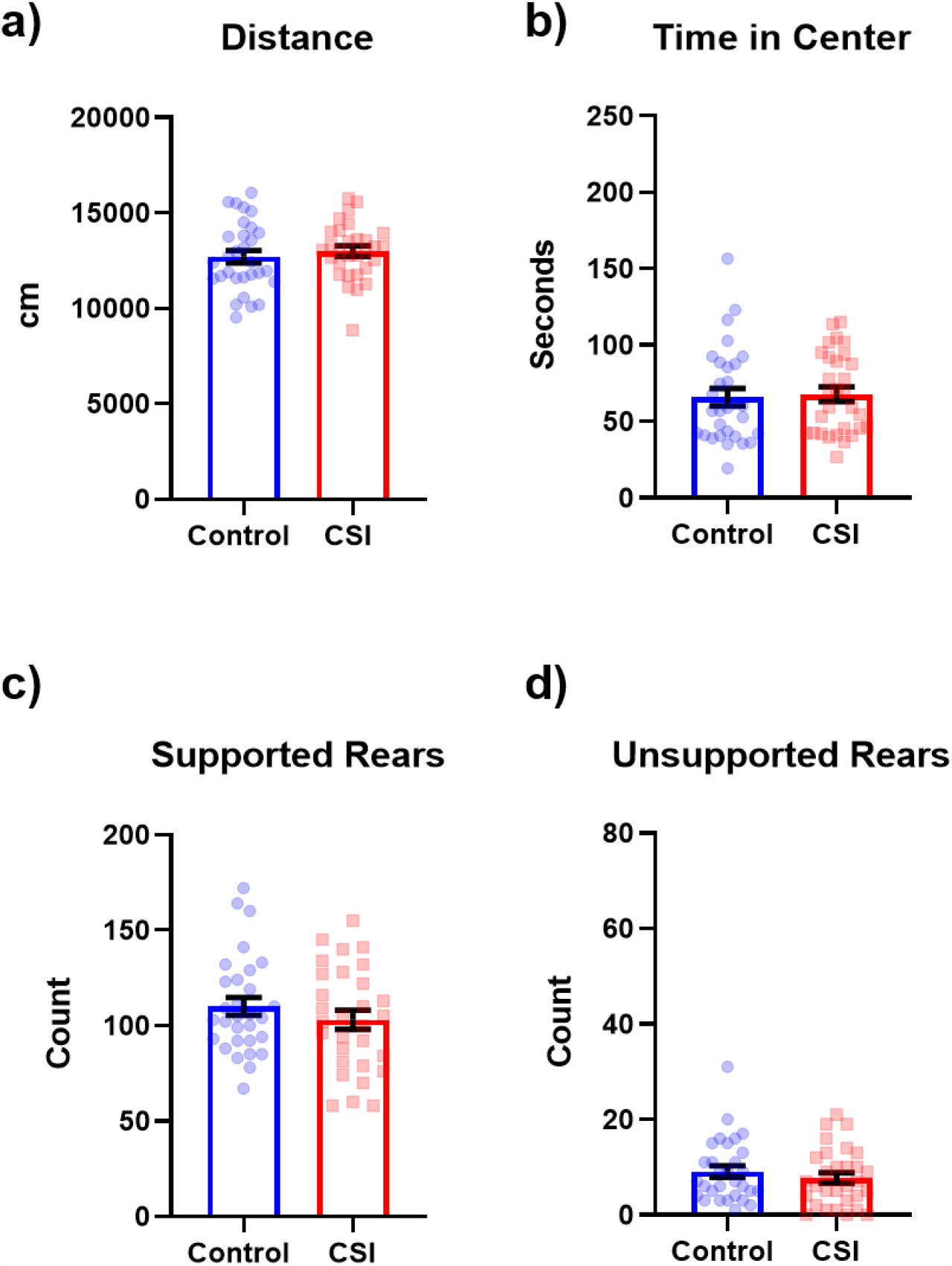
Behavioural performance of all mice before exposure to CSI stress. **(a-d)** Distance, time in center, supported and unsupported rears in the open field test in cohort 1 prior to CSI exposure as reported by the TSE Multi Conditioning system. Data expressed as mean ± SEM. Control (n=30) CSI (n=30).

**Supplementary Figure 2.**
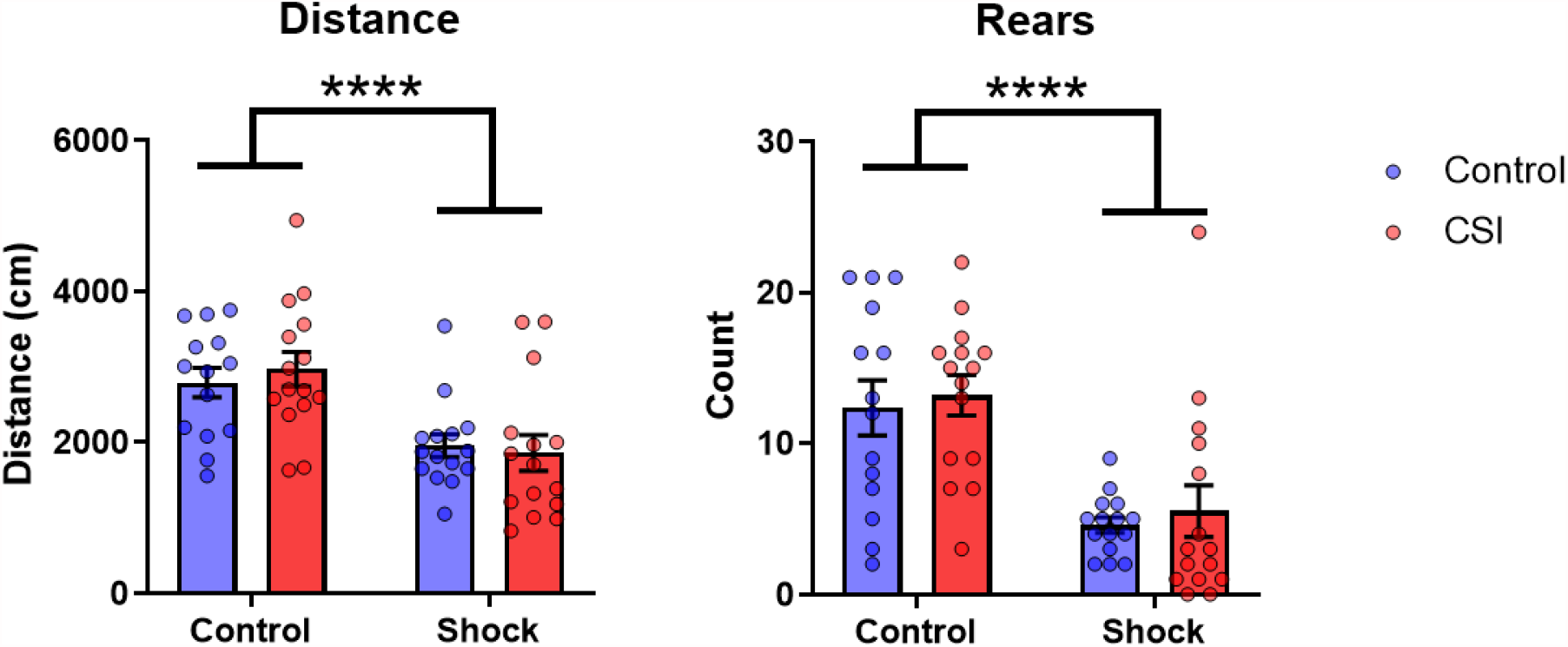
Behaviour of CSI and control mice upon re-exposure to the shock context. **(a)** Distance and **(b)** rears as reported by the TSE Multi Conditioning system in cohort 1. Data expressed as mean ± SEM. Control (n=29) CSI (n=30).

**Supplementary Figure 3.**
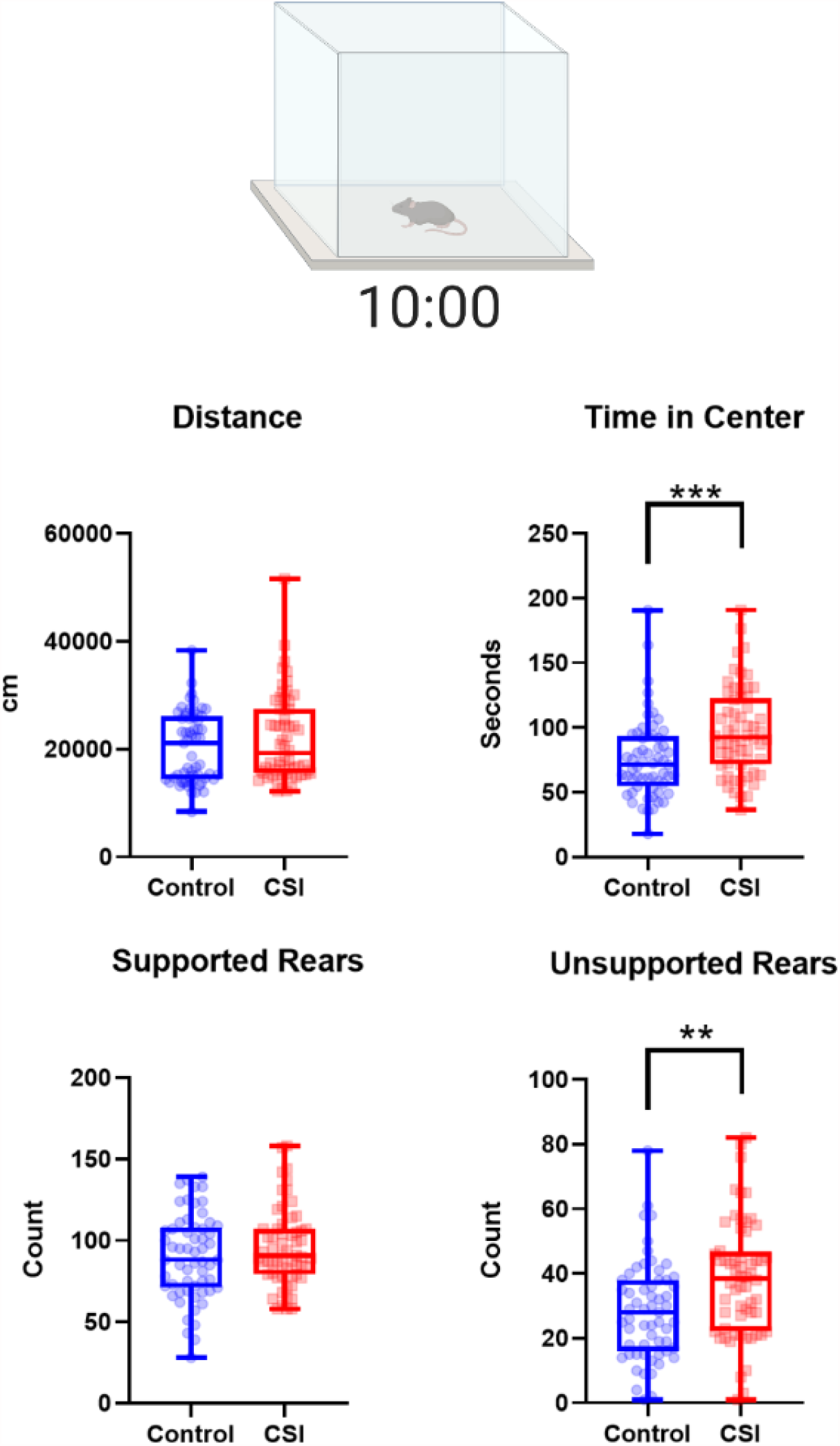
Pooled open field test data from cohorts 1 and 2. (**a-d)** Data from Figures 1 and 5 reanalyzed with the TSE Multi Conditioning system for both cohorts to make the data directly comparable. **=p<0.01,***=p<0.001.

